# MCLand: A Python program for drawing emerging shapes of Waddington’s epigenetic landscape by Monte Carlo simulations

**DOI:** 10.1101/2024.01.15.575795

**Authors:** Ket Hing Chong, Xiaomeng Zhang, Zhu Lin, Jie Zheng

## Abstract

Waddington’s epigenetic landscape is a powerful metaphor for illustrating the process of cell differentiation. Recently, it has been used to model cancer progression and stem cell reprogramming. User-friendly software for landscape quantification and visualization is needed to allow more modeling researchers to benefit from this theory.

**Results:** We present MCLand, a Python program for plotting Waddington’s epigenetic landscape with a user-friendly graphical user interface. It models gene regulatory network (GRN) in ordinary differential equations (ODEs), and uses a Monte Carlo method to estimate the probability distribution of cell states from simulated time-course trajectories to quantify the landscape. Monte Carlo method has been tested on a few GRN models with biologically meaningful results. MCLand shows better intermediate details of kinetic path in Waddington’s landscape compared to the state-of-the-art software Netland.

**Availability and implementation:** The source code and user manual of MCLand can be downloaded from https://mcland-ntu.github.io/MCLand/index.html.

## 1. Introduction

Waddington’s epigenetic landscape has been widely accepted as a metaphor for depicting cell differentiation during development^1,2^. Waddington proposed that embroynic development is hypothesized as balls rolling down different pathways of a landscape with ridges and valleys. After some times, the balls then finally reached different attractors which correspond to different stable cell states. The Waddington’s epigenetic landscape is governed by gene regulatory networks (GRN). The ground-breaking experiments demonstrating stable differentiated cells can be reprogrammed into embryonic-like stem cells^3,4,5^ have stimulated growing interest in computational modeling of the Waddington’s landscape. Recently, the Waddington’s landscape has been used to model and visualize diseases and stem cell reprogramming^6,7,8,9,10,11^.

There are many computational softwares or packages for modeling biological systems such as XPPAUT^12^, COPASI^13^, Systems Biology Toolbox for MATLAB^14^ and MATLAB solvers that provide numerical solution for time-course simulation. However, they do not support the modeling and visualization of Waddington’s epigenetic landscape. Our group has developed a software tool called NetLand^15^ for plotting the landscape based on self-consistent mean field approximation^7^. However, NetLand is not able to give details of the landscape shape such as the kinetic path between two attractors. Recently we proposed a Monte Carlo method to plot and visualize Waddington’s epigenetic landscape ^16^. In this paper, we constructed a Python program called MCLand that implement the Monte Carlo method to draw Waddington’s epigenetic landscape. MCLand can draw spontaneously emerging kinetic paths and intermediate details in Waddington’s epigenetic landscape and the results in MCLand are biologically meaningful as shown in Zhang et. al’ s work^16^.

## 2. Implementation and main functionalities

MCLand is implemented using Python 3 programming language and Tkinter package. The Monte Carlo algorithm proposed by Zhang *et al*. (2020) is used for calculating the quasi-potential (*U*) and plotting of Waddington’s epigenetic landscape. Hereafter, we refer to the quasi-potential as potential. The input file of MCLand is based on the file format design in `.ode` file of XPPAUT. Any models of GRN created in XPPAUT can be used in MCLand. To start the simulation, model equations and parameters contained in the `.ode` file need to be loaded (e.g. Fig. 1a). The model equations and parameter values are all the information required for calculating the time-course simulation using numerical solver `odeint` in `scipy` package. For the Monte Carlo method to works, a large number of time-course simulation starting from random initial conditions spanning the state space of the systems are required. Then the time-course data is projected to a 2-dimensional plane to obtain the probability distributions *P* (**x**) on the plane that is divided into tiny grid boxes^16^. From the formula *U* = *−*ln *P* (**x**) we calculate the potential values for the Waddington’s epigenetic landscape^18,16^.

**Figure 1:**
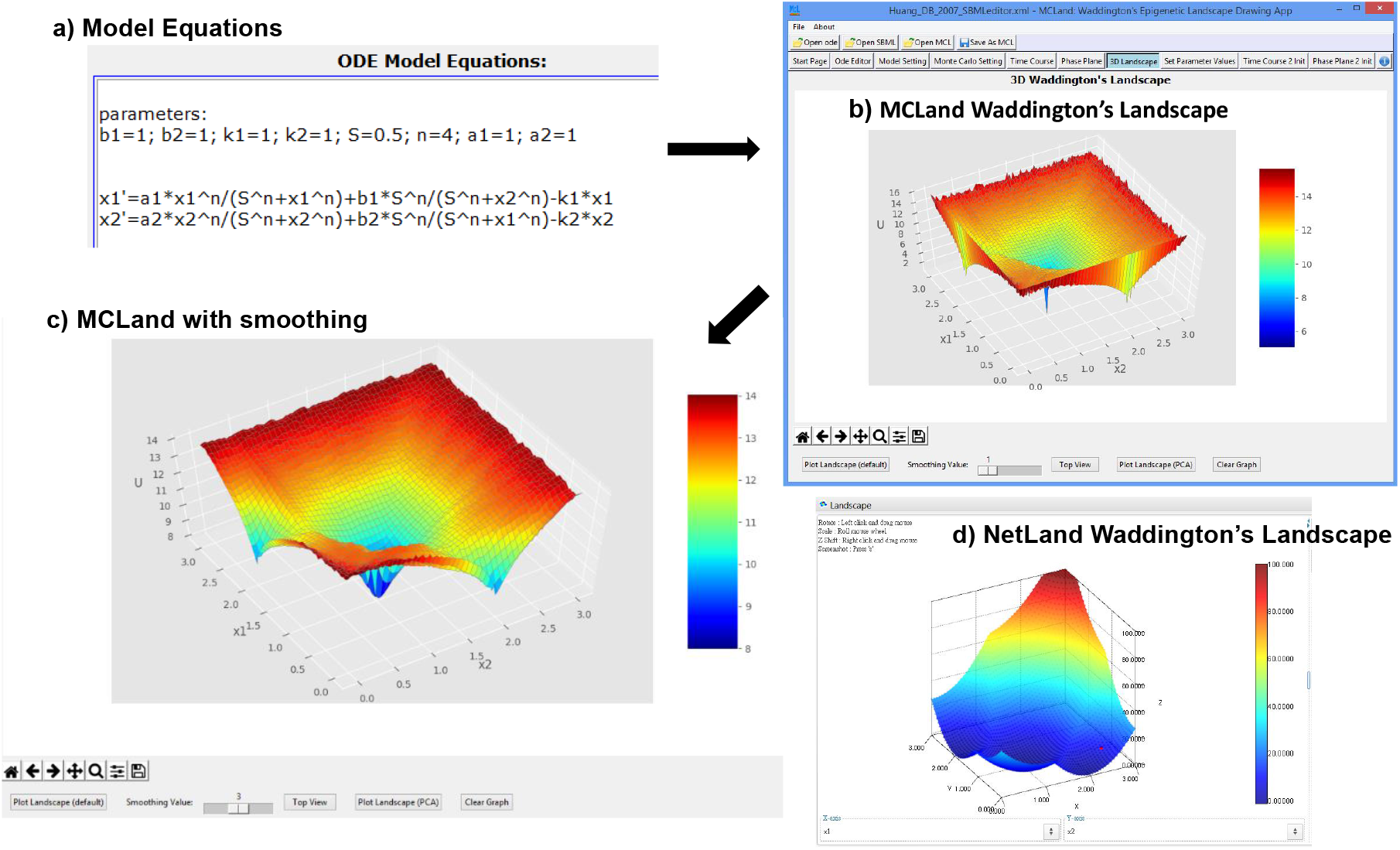
The MCLand graphical user interface result above (b) illustrates a case study from Huang et. al^17^ compared to the result from NetLand (d), all landscapes are generated using 50,000 time-course trajectories. The model of multipotent progenitor cells displays three attractors for three different cell states of progenitor, erythroid and myeloid. (a) The model equation loaded from XPPAUT model file. (b) The MCLand generated Waddington’s landscape shows three attractors with emerging shape of intermediate details of kinetic paths. (c) The MCLand Waddington’s epigenetic landscape after averaging smoothing mechanism. The Waddington’s landscape with smoothing value of 3 shows better landscape shape. d) NetLand generated Waddington’s landscape display three attractors with round shape.

### 2.1 Object oriented design

The model information is stored in Python object oriented programming class declared as `Model`. Using the `Model` class we defined a conceptual `model` object that is used to represent a model of GRN. The `model` object contains many attributes for the model GRN. The key attributes are ODE equations, parameter values, Monte Carlo method settings, time-course data for plotting of one time-course simulation, probability distributions, etc. The `model` object methods include functions for calculating probability distribution and plotting of Waddington’s epigenetic landscape. The `model` object information is also used to store model simulation results in the form of MCLand file format .pckl using the `pickle` module in Python for future plotting and analysis of Waddington’s epigenetic landscape.

### 2.2 Support SBML model

MCLand also supports the Systems Biology Markup Language (SBML) model in `.xml` file format^19^. It is convenient for users to load the SBML model file into MCLand and conduct Waddington’s epigenetic landscape study. For example, users can download SBML models from BioModels database (https://www.ebi.ac.uk/biomodels/) and draw Waddington’s landscape using MCLand.

### 2.3 Main functionalities

The design of the graphical user interface (GUI) for MCLand is shown in Fig. 1b. and the main functions are describe below. The first function is plotting of Waddington’s landscape based on Monte Carlo simulations using deterministic ODEs proposed by Zhang *et al*. (2020). After a model file has been loaded into MCLand, the plotting of landscape can be done by pressing the button “Plot Landscape (default)” located at the bottom left of Fig. 1b. The method proposed by Zhang *et al*. (2020) can be used to quantify global potential landscape of a GRN for high dimensional system.

Here, we give a summary of the theoretical description of the algorithm. For full derivation of the Monte Carlo method readers are refer to Zhang *et al*. (2020). First, from Chemical Master Equation (CME) defines *P* (**x**, *t*) the probability of model system being in state **x** at time *t*. The problem of finding *P* (**x**, *t*) can be solved using diffusion equations or through Monte Carlo simulations^20^. When *t* approaches to infinity or large, the probability *P* (**x**, *t*) will approach the steady-state probability *P*_*ss*_. Wang and co-workers proposed the potential *U* = *−*ln *P*_*ss*_ ^21,20^. We assume the average probability of state at **x** over the time from 0 to *T* as

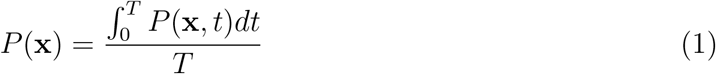

and the potential U:

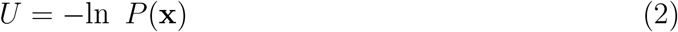

It is difficult to solve analytically for *U* = *−*ln *P* (**x**). Here, we use Monte Carlo simulations by using a large number of time-course trajectories from random initial conditions and then estimate the probability distribution of *P* (**x**). Next, we use discretize formulation to estimate the probability *P* (**x**). The time-course trajectories from random initial conditions can be simulated using Gillespie’s algorithm. However, the computational cost for Gillespie’s algorithm to get the time-evolution trajectories of CME is very high. When the number of molecules present in the biochemical reactions is large, the results from stochastic simulation is almost equivalent to deterministic simulation^22^. So, we can choose to use deterministic ODEs to speed up the calculations. Finally, we apply coarse graining step of projecting the time-course trajectories of two selected dimensions (or proteins) into a 2D plane that is divided to grid boxes. To plot landscape for a dynamical systems of high-dimensional state spaces is difficult and therefore we use a simple method of projecting trajectories into 2-dimensional plane. We can then obtain an estimate of the probability distribution *P* (**x**) and then calculate the potential U as in equation (2). An example of the potential landscape calculated using Monte Carlo method implemented in MCLand is shown in Fig. 1b. Indeed, in Zhang et. al(2020)^16^ we have shown the Waddington’s landscape obtained using stochastic simulation of Chemical Langevin Equation is equivalent to the landscape by using deterministic simulation of ODEs.

The second function is Waddington’s landscape plotting with smoothing effect. One weakness of Monte Carlo method is the landscape produced contains rugged surface. In MCLand, We proposed a simple averaging algorithm to plot smoother landscape surface. Smoothing of rugged surface of landscape can be done by changing the smoothing value of a sliding bar (next to the “Plot Landscape (default)” button). Smoothing value of 1 means no smoothing effect. For example, with smoothing value of 3 gives a good smoothing effect on the Waddington’s landscape as shown in Fig. 1c. The first and second functions mentioned above can be very useful for modelers to plot their Waddington’s landscape and gain insights for their models of GRN such as in stem cell regulations.

The third function is plotting of Waddington’s landscape with dimensionality reduction using PCA. In Zhang *et al*. (2020) we tested the Monte Carlo method in plotting of Waddington’s landscape of high dimensional system with PCA instead of two selected proteins. The plotting of MCLand with PCA can be done by pressing the “Plot Landscape (PCA)” button. The Waddington’s landscape is plotted by using the principal component 1 and the principal component 2 from PCA as the x and y axes of the landscape. We have shown that the number of attractors are consistent with the one obtained using two selected proteins^16^. For other information on how to use MCLand please refer to the user manual provided in the supplementary information.

### 2.4 A case study comparing the results from MCLand and NetLand

We conducted a case study comparing the Waddington’s landscape drawn using MCLand and NetLand. A MCLand Waddington’s landscape for the model from Huang et al. (2007) is given in Fig. 1b. MCLand can plot Waddington’s epigenetic landscape with intermediate details of kinetic paths. In the plot of NetLand (see Fig. 1d) that uses self-consistent mean field approximation method, it applies one Gaussian distribution localized at one attractor and a weighted sum of all the probability distributions for the total probability of a multistability system^7,9^. The landscape plotted by NetLand shows attractors look round and smooth, and thus it could lose some intermediate details (rugged surface, spontaneously emerging paths) which could be biologically meaningful. However, one weakness in MCLand is the rugged landscape surface as mentioned earlier. To address this problem we propose an averaging mechanism to plot a smoother landscape surface (see Fig. 1c). The MCLand Waddington’s landscape looks better after smoothing effect. Comparing Waddington’s landscape plotted between MCLand with smoothing (Fig. 1c) and NetLand (Fig. 1d), we can see MCLand with smoothing display a better quality landscape. Moreover, Monte Carlo method has been tested on a few GRN models with biologically meaningful results^16^.

## 3. Conclusion

The software NetLand can plot landscape with round and smooth attractors, but it fails to capture the details of landscape shape. Compared to the state-of-the-art software NetLand, MCLand can plot Waddington’s epigenetic landscape with intermediate details of kinetic paths. The shape of landscape in MCLand with kinetic paths reflects some biological mechanism of stability which is missing in NetLand. With smoothing effect, MCLand can produce better Waddington’s landscape shape. Overall, MCLand has a slight advantage over NetLand in terms of the quality of the Waddington’s landscape.

## Supplementary information

A user manual is available at https://mcland-ntu.github.io/MCLand/index.html

## Acknowledgements

The authors would like to thank Harrison Kinsley for providing examples of Python tkinter GUI usage. We also thank Frank Bergmann for the sharing of his Python 2 code for reading SBML model and converting it into ODEs for numerical time course simulation.

